# Species-dependent accumulation of PsaA in etioplasts points to light-independent steps in Photosystem I biogenesis

**DOI:** 10.64898/2026.06.25.734457

**Authors:** Anna Węgrzyn, Krzysztof Wardak, Radosław Mazur, Kinga Gołębiewska, Piotr Gawroński, Łucja Kowalewska

## Abstract

Whether Photosystem I (PSI) core subunits accumulate prior to light exposure in developing angiosperm seedlings remains unresolved, with conflicting reports across species. Here, we investigated the presence and membrane colocalization of the PSI core subunit PsaA in etioplasts of dark-grown angiosperms representing dicot and monocot species. Immunoblotting showed that PsaA accumulates in etioplasts of all three dicot species examined (pea, Arabidopsis, and runner bean), whereas in the monocot oat it was detected only after prolonged etiolation, at substantially lower levels and with an anomalously high apparent molecular weight. Blue-native PAGE analysis reveals that a fraction of PsaA co-migrates with LPOR, PsaB, FNR, and chlorophyll synthase, suggesting co-localization within a shared membrane microdomain rather than stable complex formation. The thylakoid insertase Alb3 was more abundant in dicot etioplasts, consistent with a potential role in the early integration of PsaA into the membrane. Upon illumination, pea reached PSI functionality faster than oat, with P700 oxidation detectable 30 min earlier, linking the dark accumulation of PsaA to an accelerated photosynthetic onset. These findings demonstrate light-independent accumulation of a PSI core subunit in a species-dependent manner and point to early steps in PSI biogenesis that precede full photosynthetic complex assembly.

**Highlight:** Contrary to prevailing models, a Photosystem I core subunit PsaA accumulates in dark-grown angiosperm seedlings before light exposure, revealing light-independent early steps in photosynthetic complex biogenesis.

## Introduction

The thylakoid membranes of chloroplasts house the multiprotein complexes responsible for the light-dependent reactions of photosynthesis, including Photosystems I and II (PSI and PSII), which absorb light and drive electron transport via a chain of redox reactions (Johnson, 2016; Johnson, 2025). PSI plays a particularly critical role downstream, receiving electrons from plastocyanin and transferring them to ferredoxin (Schöttler *et al*., 2011).

PSI is composed of both chloroplast-encoded (PsaA, PsaB, PsaC, PsaI, PsaJ) and nuclear-encoded (PsaD, PsaE, PsaF, PsaG, PsaH, PsaK) subunits, making its biogenesis particularly intricate in plants (Mahapatra *et al*., 2024; Naschberger *et al*., 2024; Schöttler *et al*., 2011). Although the structure and function of PSI have been extensively studied (Naschberger *et al*., 2024; Schöttler *et al*., 2011), the sequence of events governing the assembly of its core components remains incompletely understood (Rolo *et al*., 2024a; Wittenberg *et al*., 2017). This is largely due to the rapid pace of PSI assembly (Kim *et al*., 2024; Zhang *et al*., 2024), which must be tightly coordinated both spatially and temporally (Rolo *et al*., 2024a). Assembly predominantly occurs in young, developing leaf tissue, after which PSI forms a stable, low-turnover complex, making isolation of assembly intermediates particularly challenging (Rolo *et al*., 2024a).

PSI assembly is initiated by the cotranslational insertion of the chloroplast-encoded subunits PsaA and PsaB into the thylakoid membrane, forming the reaction center heterodimer (Schöttler *et al*., 2011). However, in higher plants, the lack of structural data on intermediate forms has limited our understanding of this process. Recent research on oat (*Avena sativa*) has identified a native subcomplex, pre-PSI-1, shedding light on conserved steps in PSI biogenesis (Naschberger *et al*., 2024). Another intermediate, PSI*, comprises a specific subset of PSI core subunits (PsaA, PsaB, PsaC, PsaD, PsaE, PsaH, PsaI, and PsaL), but lacks the light-harvesting complex I (LHCI) antenna proteins. PSI* accumulates in high amounts in young leaf tissue – where active thylakoid formation takes place – and its abundance decreases with leaf maturation, indicating its role as a transient assembly intermediate (Wittenberg *et al*., 2017). Its identification provides a valuable biochemical entry point for studying PSI biogenesis and the roles of various assembly factors in thylakoid development.

In angiosperms, given the complexity of PSI assembly, numerous chaperones and auxiliary factors are involved. Albino3 (Alb3) facilitates the membrane insertion of PsaA and PsaB (Göhre *et al*., 2006), working in concert with PPD1 (Liu *et al*., 2012), the plastid-encoded Ycf3 (Albus *et al*., 2010; Naver *et al*., 2001; Ruf *et al*., 1997), and the Ycf4, which specifically interacts with nascent PSI subunits (Naver *et al*., 2001; Nellaepalli *et al*., 2018; Onishi and Takahashi, 2009). Translational regulation plays a central role in determining the accumulation of chloroplast-encoded PSI subunits (Dall’Osto *et al*., 2013; Lezhneva and Meurer, 2004). Additional regulatory mechanisms have been identified that affect PSI biogenesis. The PBR1–Ycf1 module has been described as a translational control element for PSI and other photosynthetic complexes (Yang *et al*., 2016), and HCF145 has been implicated in the transcriptional regulation of the psaA-psaB-rps14 operon (Lezhneva and Meurer, 2004). Moreover, studies in Chlamydomonas (*Chlamydomonas reinhardii*) have proposed a hierarchical model of autoregulation in which the synthesis of PsaA and PsaC depends on the presence of PsaB, without which translation is halted (Rochaix, 2011; Wostrikoff *et al*., 2004). The prevailing model suggests that PsaA synthesis requires (i) light-dependent activation, (ii) de novo chlorophyll biosynthesis, and (iii) the prior presence of PsaB (Chitnis, 2001; Eichacker *et al*., 1990; Wostrikoff *et al*., 2004). However, this model is mainly supported by studies on mature chloroplasts (Baker and Leech, 1977; Stöckel and Oelmüller, 2004; Wang *et al*., 2020) and unicellular organisms (Jin *et al*., 2008; Sun *et al*., 2019; Zhang *et al*., 2014), and whether it applies during plant ontogenesis remains to be clarified (Yang *et al*., 2015). Moreover, recent studies indicate that this hierarchical translational feedback is not strictly conserved in land plants, where the synthesis of PSI core subunits appears to be less tightly coupled to complete PSI assembly than in Chlamydomonas (Ghandour *et al*., 2025), indicating a potential divergence in regulatory mechanisms between algae and higher plants. Furthermore, the discovery of the land-plant-specific assembly factor PBF8 revealed previously unknown PSI assembly intermediates: PSI core subcomplex I, containing PsaA–E, PsaH, PsaI and PsaL, and PSI core subcomplex II, formed upon the incorporation of PsaF. This finding suggests that additional PSI biogenesis mechanisms evolved during plant terrestrialization, highlighting further divergence between algal and land-plant PSI assembly pathways (Zhang *et al*., 2024).

PSI biogenesis in dark-grown (etiolated) angiosperm seedlings remains underexplored. Some studies report the presence of PsaA in etioplasts of Arabidopsis (*Arabidopsis thaliana*), bean (*Phaseolus vulgaris*), oat, pea (*Lathyrus oleraceus*), and in cotyledons of transgenic Arabidopsis seedlings expressing 35S::PBR1-YFP (Li *et al*., 2020; Nechushtai and Nelson, 1985; Rudowska *et al*., 2012; Yang *et al*., 2016), while others have not detected it in barley (*Hordeum vulgare*), pea, and common wheat (*Triticum aestivum*) (Eichacker *et al*., 1990; Kanervo *et al*., 2008; Klein *et al*., 1988; Laing *et al*., 1988; Machold and Høyer-Hansen, 1976; Takabe *et al*., 1986; Vierling and Alberte, 1983). Notably, positive detections have been reported predominantly in dicot species, whereas most etioplast proteomes lacking PsaA were derived from monocots, raising the question of whether these discrepancies reflect methodological limitations or genuine species-specific differences in PSI core protein accumulation. In line with these observations, several other PSI subunits appear to be present in etioplasts prior to illumination (Li *et al*., 2020; Nechushtai and Nelson, 1985; Rudowska *et al*., 2012; Takabe *et al*., 1986; Yang *et al*., 2016), though their abundance may vary across species. Nonetheless, proteomic analyses have identified some components associated with PSI assembly in etioplasts, such as Ycf4 in barley (Plöscher *et al*., 2011) and a PSI assembly-related stress-inducible protein in rice (*Oryza sativa*) (Blomqvist *et al*., 2008), suggesting that elements of the assembly machinery may be present even in the absence of fully assembled PSI.

Etiolated seedlings exhibit characteristic adaptations aimed at maximizing their chances of reaching light and transitioning to autotrophic growth. Their plastids, termed etioplasts, contain a diamond-type cubic prolamellar body (PLB) connected to lamellar prothylakoids, which serve as precursors to the thylakoid membranes of mature chloroplasts. Although the organization of these membranes in etioplasts differs from that of chloroplasts, they may already provide a spatial framework for the early stages of photosynthetic protein biogenesis. Supporting this idea, recent studies in Chlamydomonas have shown that distinct membrane domains play key roles in the translation and stepwise assembly of membrane proteins, within a spatially and temporally coordinated system (Sun *et al*., 2025).

Upon light exposure, de-etiolation is initiated, involving the reorganization of internal membranes into grana and stroma thylakoids, as seen in functional chloroplasts (Armarego-Marriott *et al*., 2020). This process is accompanied by a rapid upregulation of photosynthetic protein and pigment synthesis (Armarego-Marriott *et al*., 2019) and requires tight coordination between nuclear and plastid gene expression, regulated at multiple levels, including light-responsive signaling pathways (Kafri *et al*., 2023; Mahapatra *et al*., 2024). A hallmark event is the light-dependent reduction of protochlorophyllide (Pchlide) to chlorophyllide by light-dependent Pchlide:NADPH oxidoreductase (LPOR - localized in PLB), facilitating the conversion essential for completing chlorophyll biosynthesis and enabling the formation of pigment–protein complexes essential for efficient energy capture (Adam *et al*., 2011; Floris and Kühlbrandt, 2021; Solymosi and Schoefs, 2010). Although both photosystems begin to assemble during seedling greening, PSI activity has been reported to precede that of PSII (Krol *et al*., 1987; Ohashi *et al*., 1989), suggesting that PSI biogenesis may initiate or progress more rapidly in the early stages of de-etiolation.

During de-etiolation, light triggers the accumulation of PSI core subunits PsaA and PsaB, along with chlorophyll (Høyer-Hansen and Simpson, 1977; Klein and Mullet, 1986; Nechushtai and Nelson, 1985; Vierling and Alberte, 1983). However, the molecular events preceding light exposure remain poorly understood. The *psaA* and *psaB* genes are arranged in tandem and co-transcribed (Hihara and Sonoike, 2001; Kirsch *et al*., 1986), with mRNA levels in etioplasts comparable to those in chloroplasts (Ji *et al*., 2022; Klein and Mullet, 1986; Kreuz *et al*., 1986; Laing *et al*., 1988; Sandoval-Ibáñez *et al*., 2021). In etiolated seedlings, these transcripts are found on membrane-bound polysomes (Hihara and Sonoike, 2001; Klein *et al*., 1988; Kreuz *et al*., 1986; Laing *et al*., 1988), yet PsaA translation appears to depend on the prior presence of PsaB (Hihara and Sonoike, 2001; Stampacchia *et al*., 1997), suggesting a regulatory mechanism that operates despite co-transcription. Their translation is thought to be primarily controlled at the post-transcriptional level, limiting the accumulation of PSI core proteins in the absence of light (Bai *et al*., 2017; Hihara and Sonoike, 2001; Ji *et al*., 2022; Klein and Mullet, 1986; Kreuz *et al*., 1986; Laing *et al*., 1988). Nonetheless, in vitro studies show that full-length PsaA and PsaB can be synthesized in etioplast polysomes even in the absence of light (Klein *et al*., 1988).

Emerging evidence suggests that the chloroplast redox state and NAD kinase (NADK) activity influence the translation of PsaA and PsaB during PSI biogenesis (Bhattacharjee, 2013; Ji *et al*., 2022), underscoring the intricate interplay between metabolic signals and protein synthesis in photosynthetic development. Despite these advances, the regulatory mechanisms governing PsaA and PsaB expression in dark-grown angiosperms remain poorly understood and warrant further investigation (Hihara and Sonoike, 2001). However, even the more basic question of whether PSI core subunits accumulate in etioplasts prior to light exposure remains unresolved. While psaA and psaB transcripts are consistently detected in darkness, data on their translation are conflicting. We hypothesize that a pool of PSI apoproteins accumulates during etiolation, establishing molecular prerequisites for PSI assembly prior to light exposure. This study aims to resolve this inconsistency by comparing PSI core protein accumulation across monocot and dicot species and examining the co-localization of PsaA with other proteins in etioplast membranes. Moreover, we investigate the relationship between PsaA dark accumulation and PSI functional readiness in the early stages of chloroplast biogenesis.

## Materials and Methods

### Plant species and growing conditions

*Lathyrus oleraceus* (purchased as *Pisum sativum* L. cv. Iłówiecki; PNOS, 05-850 Ożarów Mazowiecki, Poland), *Avena sativa* (PNOS, 05-850 Ożarów Mazowiecki, Poland), *Phaseolus coccineus* L. cv. Eureka (PlantiCo Zielonki, 05–082 Babice Stare, Poland), and *Arabidopsis thaliana* (ecotype Columbia-0, N1092) were selected for experimental procedures. Embryos (0 days of etiolation) and first leaves or cotyledons for Arabidopsis (3, 5, 7, and 10 days of etiolation) were collected for various experiments. Ten-day-etiolated seedlings were exposed to PAR light (150 µmol photons m⁻² s⁻¹) in a climate-controlled room for 3 h. As controls, 20-day-old plants grown under a 16 h light/8 h dark photoperiod (21 °C day/20 °C night) at the same PAR intensity in perlite-containing pots were used; fully expanded leaves were collected for analysis. Plants were fertilized with modified Knop’s nutrient solution.

All procedures involving etiolated plants were conducted under strict dark conditions, using only dim, indirect green light as the sole light source.

### Preparation of plastids and protein extracts

Approximately 100–200 mg of plant material, previously frozen in liquid nitrogen, was ground to a fine powder in a pre-chilled mortar and transferred to extraction buffer (50 mM Tris-HCl, pH 8.0; 10% (v/v) glycerol; 2% (w/v) SDS; 25 mM EDTA; 1 mM PMSF). Samples were thoroughly mixed, frozen in liquid nitrogen, and subjected to three freeze–thaw cycles using a 4 °C ultrasonic bath. After the final thaw, extracts were centrifuged at 10,000 × g, and the resulting supernatants were collected for further analysis (Krysiak *et al*., 2024).

Chloroplasts from mature plants were isolated by homogenization in a buffered isotonic medium and subsequent filtration through 4 layers of Miracloth (Ø 50 μm) (Merck Millipore) and centrifugation as described in (Krysiak *et al*., 2024). The chlorophyll concentration was quantified spectrophotometrically after extraction with 80% (v/v) acetone (Lichtenthaler, 1987).

To isolate etioplasts, first leaves of etiolated seedlings were homogenized (Waring blender) in buffer containing 0.5 M sucrose, 20 mM Tricine, 10 mM Hepes, 50 mM KCl, 5.5 mM cysteine, 10 mM NaF, 1 mM PMSF, and 15 μM NADPH, adjusted to pH 8.0. The homogenate was filtered through four layers of Miracloth and centrifuged at 3,300 × g for 15 min at 4 °C. The pellet was resuspended in the same buffer and centrifuged again at 4,340 × g for 10 min at 4 °C. The final pellet contains etioplast crude fraction.

### Low-Temperature fluorescence measurements

Low-temperature (77 K) fluorescence emission spectra were recorded using a modified Shimadzu RF5301-PC fluorometer system. Plant material (∼20 mm²) was placed in a PTFE cuvette on a quartz holder connected to the system via optical fibers. Samples were frozen in liquid nitrogen and excited at 440 nm (excitation slit 10 nm). Emission spectra were recorded in the range 600-800 nm (emission slit 5 nm) with a 1 nm interval through the LP600 long-pass filter. Raw spectra were background-corrected and normalized as specified in the figure captions.

### SDS-PAGE and Western blot

Samples containing 10 μg of protein or 1-3 μg of chlorophyll were denatured in Laemmli buffer, loaded onto a polyacrylamide gel, and separated using the standard SDS-PAGE protocol (Krysiak *et al*., 2024). SDS-PAGE–resolved proteins were either stained with Coomassie Brilliant Blue (R-250) or electrotransferred to a PVDF membrane. Transfer was performed at a constant voltage of 100 V for 45 min at room temperature using Trans-Blot wet transfer system (Bio-Rad). Western blotting was performed as described previously (Krysiak *et al*., 2024)) using primary and secondary antibodies (Agrisera, Vännäs, Sweden) as follows: PsaA (AS06 172), PsaB (AS10 695), PsaC (AS10 939), PsaD (AS09 461), Lhca2 (AS01 006), POR (AS05 067), FNR (AS15 2909), ChlG (AS14 2793), Alb3 (AS24 5026), and Ycf4 (AS24 5027), HRP-conjugated anti-rabbit IgG (AS09 602).

### Blue Native PAGE

Plastid samples were treated according to (Bykowski *et al*., 2021)) except that solubilization was performed using n-dodecyl-β-D-maltoside and digitonin, each to a final concentration of 1% (w/v), followed by 30 min incubation at 4 °C in the darkness with agitation (Sárvári *et al*., 2022). Samples from green tissue were normalized to 8.3 µg chlorophyll per lane. For etiolated seedlings, which lack chlorophyll, the total protein amount loaded was matched to the protein content present in green samples at 8.3 µg chlorophyll. The prepared samples were loaded onto NativePAGE™ 4–16%, Bis-Tris 1.0 mm thick gradient gels (Invitrogen), and protein complexes were resolved as described in (Bykowski *et al*., 2021)).

Resolved BN-PAGE gels were incubated at 4 °C for 20 min with gentle shaking (40 RPM) in a 300 mM tris-acetate (pH 8.6) buffer containing 100 mM acetic acid, and 1% (w/v) SDS, then transferred to fresh buffer and enclosed between two glass plates for 40 min at 4 °C without agitation. Protein complexes were transferred onto PVDF membrane (0.2 μm, Bio-Rad) using the Trans-Blot wet transfer system (Bio-Rad) at a constant current of 60 mA (15 V limit) for 16 h at 4 °C with continuous cooling and stirring in transfer buffer (150 mM Tris-acetate (pH 8.6), 50 mM acetic acid) (modified from (Wu *et al*., 2021). After transfer membranes were washed (3 x 1 min) with methanol at RT, followed by two 5-minute washes in Tris Buffered Saline solution. Western blotting was then performed as described above.

### RNA extraction and qRT-PCR analysis

Total RNA was isolated from 25–50 mg of tissue using a home-made TRIzol substitute, following a standard protocol (Rodríguez-Ezpeleta *et al*., 2009). Subsequently, 10 µg of RNA was treated with DNase I (New England Biolabs, M0303) and purified by sodium acetate/isopropanol precipitation. The cDNA was synthesized from 500 ng of RNA using the High-Capacity cDNA Reverse Transcription Kit (Applied Biosystems, 4368814), according to the manufacturer’s instructions, in a scaled-down 10 µL reaction. Quantitative real-time PCR (RT-qPCR) was performed in 10 µL reaction volumes containing 1× Luna Universal qPCR Master Mix (New England Biolabs, M3003), 0.2 µM of each primer, and 2 µL of cDNA template (corresponding to a 5-fold dilution of the cDNA reaction). Amplification was conducted on a CFX96 Real-Time PCR Detection System (Bio-Rad) using the following program: initial denaturation at 95 °C for 1 min, followed by 40 cycles of 95 °C for 15 s and 60 °C for 30 s. A melt curve analysis was then performed from 65 °C to 95 °C with 0.5 °C increments and 5 s per step. Primer sequences are listed in Supplementary Table 1. Relative gene expression levels were calculated using the ΔΔCt method, with two reference genes per species: pea – PP2A and TUBB (Die *et al*., 2010) and oat – ARF1 and HNR (Sowa *et al*., 2024). Ct values were corrected for primer efficiency as determined using LinRegPCR v2021.2.

### Protein digestion and LC-MS/MS

The etiopast samples containing 50 μg of protein were diluted to a protein concentration of 1 mg ml^-1^ in 100 mM ammonium bicarbonate containing 0.1% (w/v) RapiGest^TM^ surfactant (Waters, Milford, MA, USA). After 30 min incubation at 30 °C, 5 μg of the sequencing grade trypsin (Promega) was added, and samples were digested overnight at 30°C. After incubation, samples were centrifuged at 10,000 × g for 5 min, the supernatant was collected and supplemented with trifluoroacetic acid to a final concentration of 0.5% (v/v) following 45 min incubation at 37 °C. After centrifugation 12,000 × g for 10 min, the supernatant was filtered through a 0.22 μm tube filter and dried in a SpeedVac concentrator.

LC-MS/MS analyses were performed in the Mass Spectrometry Laboratory (Institute of Biochemistry and Biophysics, Polish Academy of Science) using an Evosep One instrument (Evosep Biosystems, Denmark) directly coupled to an Orbitrap Astral mass spectrometer (Thermo Fisher Scientific, Germany) working in Data-Independent Acquisition mode. The raw data were analyzed with DIA-NN software (version 2.1) using a reference proteome from Uniprot for *Pisum sativum* (UP001058974, version 2025_03, 1 sequence per gene, 45086 sequences) and *Arabidopsis thaliana* (UP000006548, version 2025_03, 1 sequence per gene, 27 448 sequences), and supplemented with popular contaminants. q-value threshold for results was set to 0.01.

### P700 measurements

P700 measurements were performed using a pulse-amplitude modulation fluorometer, a Dual-PAM-100 (Heinz Walz GmbH, Effeltrich, Germany). Seedlings (without dark adaptation) were mounted between the measuring heads of the device, and the PSI measuring light was balanced to achieve a signal around 0.0 V. After 10 s of far-red light illumination, a 0.3 s red light saturation pulse with 10,000 μmol photons m^−2^ s^−1^ was applied. The P700 signal was recorded from -200 ms (before the saturation pulse) to 1400 ms (after the saturation pulse), with a time resolution of 0.3 ms.

### Statistical analysis

qRT-PCR data were analyzed using the Kruskal–Wallis test followed by Dunn’s post hoc multiple-comparisons test. Differences were considered statistically significant at p < 0.05. The number of biological replicates is indicated in the figure legend.

## Results

### PSI core subunit accumulation in etioplasts across angiosperm species

To investigate whether previously reported discrepancies in PsaA detection in etiolated seedlings may reflect species-specific variation, we compared PSI subunit abundance across three dicot species: pea (*Lathyrus oleraceus*), Arabidopsis (*Arabidopsis thaliana*), runner bean (*Phaseolus coccineus*), and one monocot species: oat (*Avena sativa*). The accumulation of PsaA and PsaB in etiolated seedlings was examined relative to fully developed chloroplasts in mature plants (Fig. 1). SDS–PAGE followed by immunoblotting revealed immunospecific signals for the presence of PsaA subunit in all analyzed species. In oat, PsaA migrated with a higher apparent molecular weight compared to mature chloroplasts and etiolated dicot species, an observation that was reproducible across independent biological replicates. In contrast, PsaA in all dicot species was consistently detected at the expected molecular weight. PsaB signals were weak and close to the detection limit across all species. Among dicots, Arabidopsis exhibited a visibly stronger PsaA signal than runner bean and pea; however, as differences in antibody affinity between species cannot be excluded, these comparisons should be interpreted qualitatively. The presence of PsaA in etioplasts isolated from pea after 10 days of etiolation was confirmed by tandem mass spectrometry (MS/MS) analysis (Supplementary Table 2), independently validating the immunoblot results. The identification of PsaA-specific peptides provided proteomic confirmation of the accumulation of this PSI core subunit in the absence of light.

**Fig. 1.**
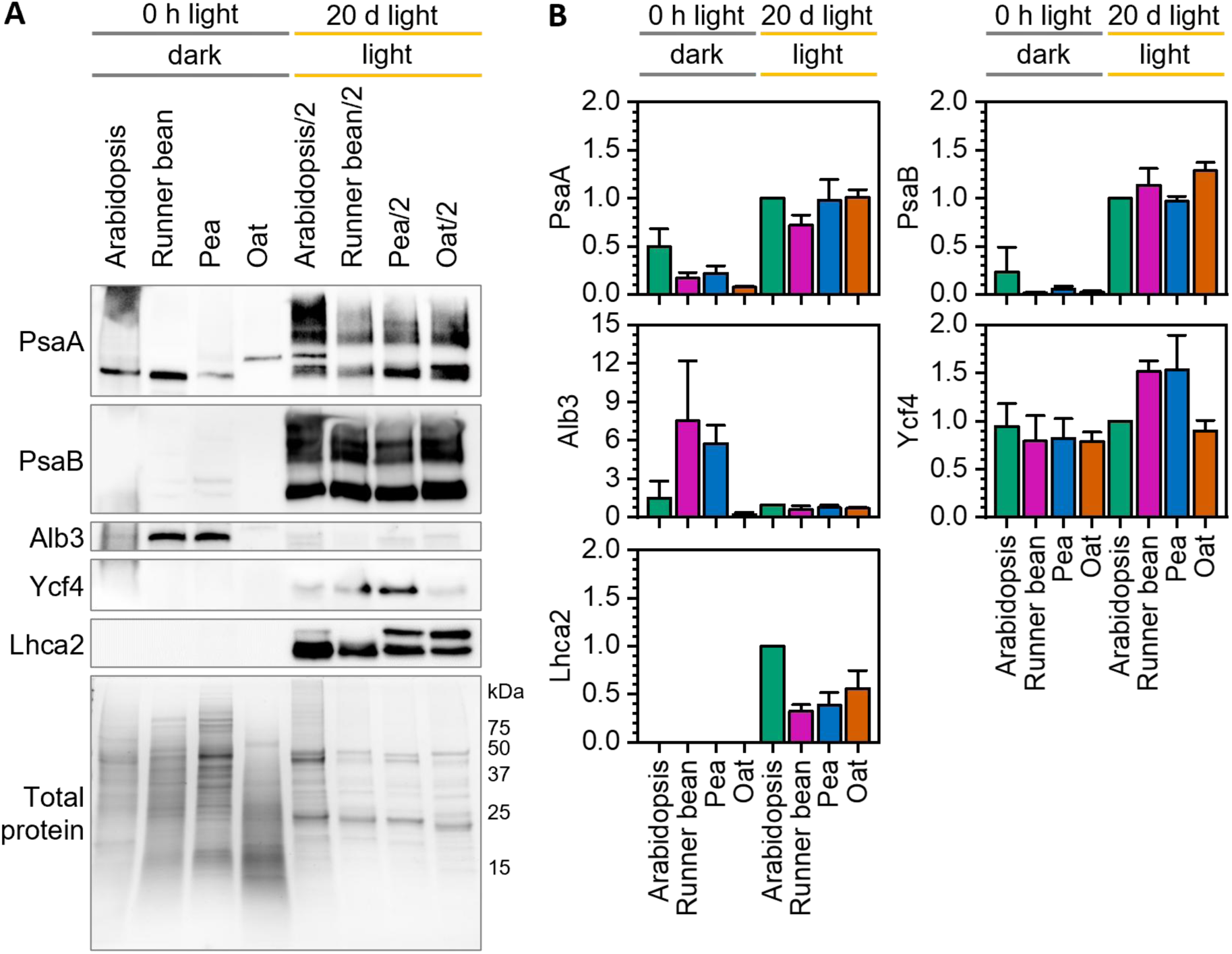
Accumulation of PSI core subunits and assembly-associated proteins in etiolated tissues of selected angiosperm species. (A) Total protein extracts from etiolated seedlings and chloroplasts isolated from mature light-grown plants of Arabidopsis (*Arabidopsis thaliana*), runner bean (*Phaseolus coccineus*), pea (*Lathyrus oleraceus*), and oat (*Avena sativa*) were separated by SDS–PAGE and analysed by immunoblotting. Proteins detected in darkness include the PSI core subunits PsaA and PsaB, the PSI assembly factor Ycf4, and the thylakoid insertase Alb3; representative blots from 3 independent biological replicates are shown. Equal loading was confirmed by Coomassie staining. (B) Signal intensities were quantified using ImageLab software version 6.0.1 (Bio-Rad) (Taylor *et al*., 2013) and normalised to the corresponding signal in mature Arabidopsis samples. Values represent means ± SD (error bars) of 3 independent biological replicates. Statistical analysis was not performed, as species-specific differences in antibody affinity limit direct quantitative comparisons between species.

We also examined two proteins associated with early PSI assembly: Ycf4 and Alb3 (Fig. 1). Ycf4 was detected in mature chloroplasts, consistent with its established role in PSI assembly in developed plastids (Plöscher *et al*., 2011). In contrast, Alb3, involved in anchoring PsaA and PsaB into the thylakoid membrane (Bellafiore *et al*., 2002; Pasch *et al*., 2005), accumulated in etioplasts, particularly in dicot species (Fig. 1). This observation supports Alb3 potential role in early stages of PSI complex formation in etiolated tissues.

### Pigment dynamics during etiolation

During etiolation, the internal membrane system undergoes progressive development, accompanied by changes in the size and organization of the PLB and prothylakoids (Bykowski *et al*., 2020). To determine whether the duration of etiolation influences the accumulation of PSI components, we monitored protein and pigment dynamics over time in oat and pea, selected for their high biomass yield. All experiments were conducted under strictly controlled dark conditions, with handling performed exclusively under dim green light to prevent unintended illumination.

Low-temperature (77 K) fluorescence spectroscopy was employed both as a control for maintaining etiolation and as a tool to monitor pigment–protein complex formation. No fluorescence signals corresponding to chlorophyll-containing complexes were detected at any time point of etiolation, confirming that seedlings remained fully etiolated throughout the experiment (Fig. 2). Fluorescence emission spectra revealed two main emission bands: ∼630 nm, corresponding to free protochlorophyllide (Pchlide), and ∼653 nm, associated with the Pchlide:LPOR:NADPH complex (Fig. 2) (Skupien *et al*., 2017). In both species, the first day of etiolation (-10d) was characterized by weak, non-specific fluorescence. By day 3 (-7d), an increase in the 653 nm signal indicated the accumulation of the Pchlide:LPOR:NADPH complex, whose contribution in the total registered signal continued to increase with prolonged etiolation, however, with species-specific dynamics.

**Fig. 2.**
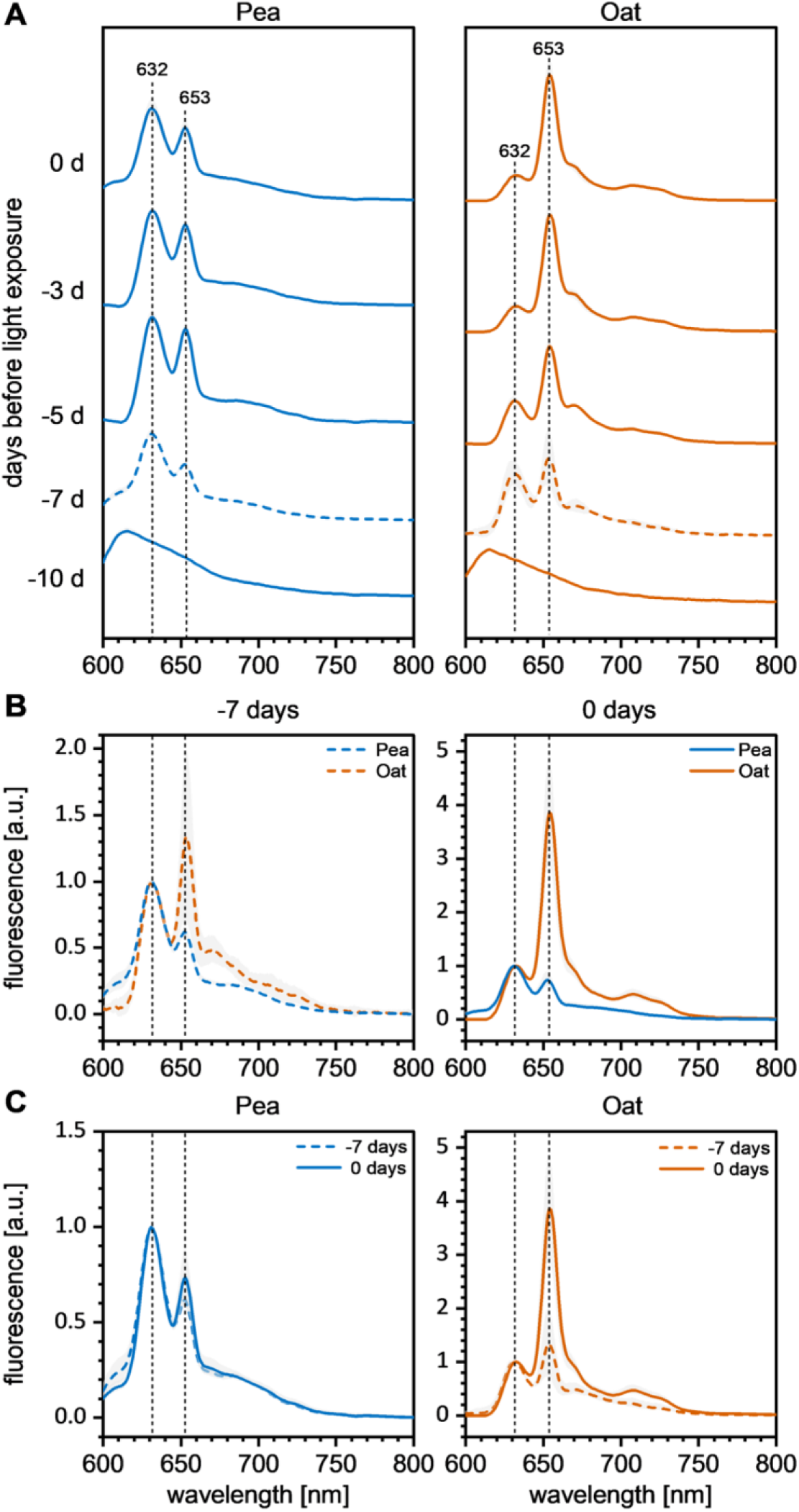
Low-temperature fluorescence spectroscopy of etiolated pea and oat seedlings. (A) Averaged 77 K fluorescence emission spectra recorded from pea and oat seedlings at successive stages of 10-day-long etiolation (−10d to 0d). Excitation wavelength: 440 nm. Emission bands at ∼632 nm and ∼653 nm correspond to free protochlorophyllide (Pchlide) and the photoactive Pchlide:LPOR:NADPH ternary complex, respectively. Spectra were normalized to the equal area under the curve. (B) Comparison between averaged spectra recorded at -7d and 0d for both species revealing changes in Pchlide form ratios. Spectra were normalized to the maximum at ∼632 nm. (C) Cross-species comparison between averaged spectra recorded at -7d and 0d. Spectra were normalized to the maximum at ∼632 nm. Shaded regions represent means ± SD of 2-6 independent biological replicates. d, “-” days of etiolation prior to illumination; e.g., -7d, 3 days of etiolation; 0d, 10-day-old etiolated seedlings.

In pea, the 653/630 nm ratio remained relatively stable throughout the experiment, ranging from 0.64 (-7d) to 0.79 (0d), indicating a steady balance between free and protein-bound Pchlide (Fig. 2). In contrast, oat showed a pronounced increase in this ratio over time, rising from 0.81 after 3 days (-7d) to 4.88 after 10 days of etiolation (0d) (Fig. 2), suggesting a more efficient accumulation of the Pchlide:LPOR:NADPH complex. Additionally, in oat, a weak fluorescence peak around 670 nm appeared from day 5 onward (-5d) (Fig. 2), consistent with the formation of free Pchlide aggregates (Skupien *et al*., 2017).

### Transcript and protein accumulation of PSI components during etiolation

To track the accumulation of PSI components during etiolation, qRT-PCR and immunoblot analyses were performed in parallel to assess transcript and protein dynamics and identify key regulatory stages in PsaA accumulation. In oat, *psaA, psaB,* and *atpβ* transcript levels showed an increasing trend during etiolation, which became significant only after 10 days (0d) compared with day 1 (-10d). In contrast, pea seedlings showed no significant changes over the same period. Upon illumination, a marked increase in *psaA*, *psaB*, and *atpβ* transcript abundance was observed in pea but not in oat, whereas *Lhca2* transcript levels increased upon illumination in both species (Fig. 3). Expression patterns of *psaA* and *psaB* were highly similar, consistent with their co-transcription (Lezhneva and Meurer, 2004).

**Fig. 3.**
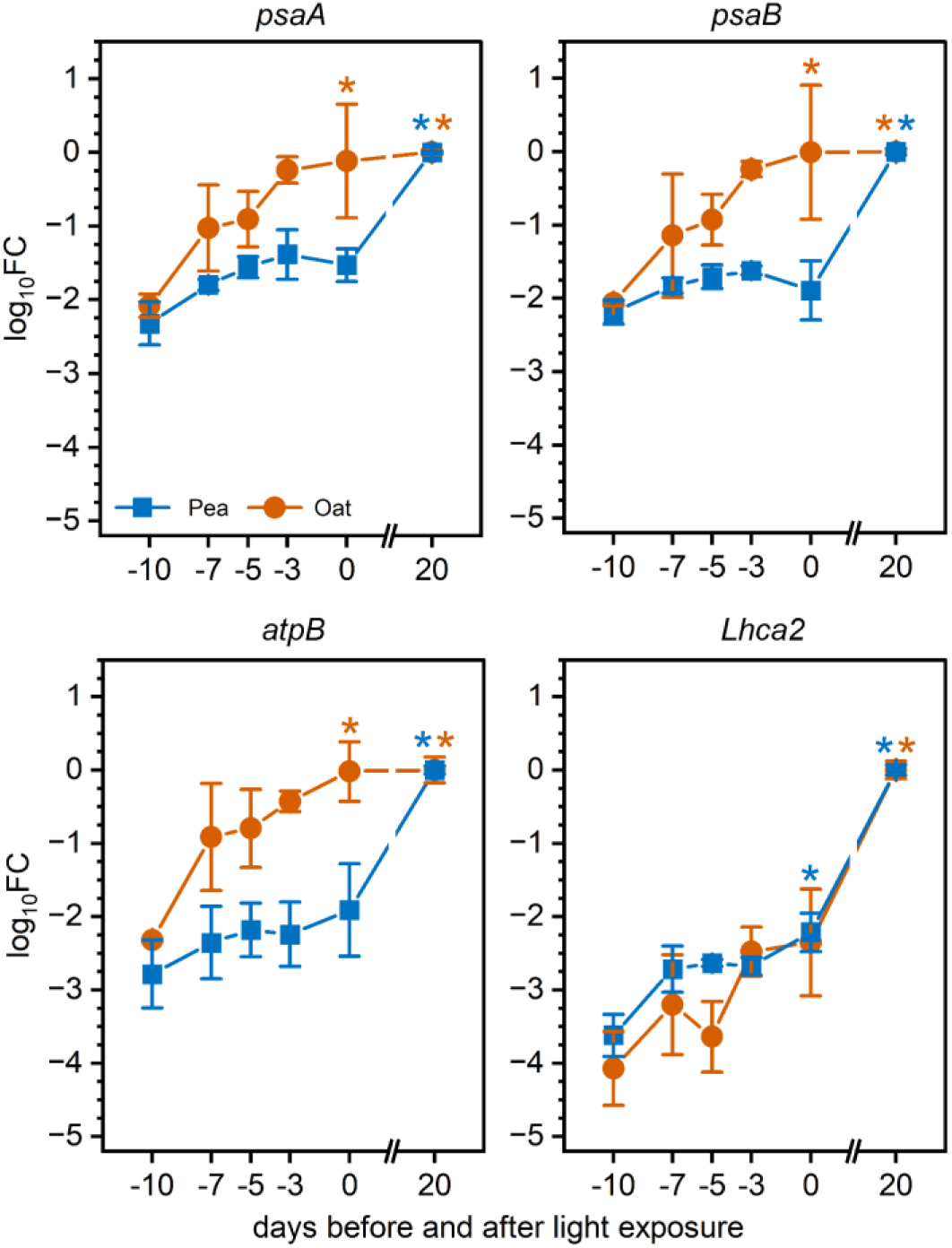
Transcript accumulation of PSI and photosynthesis-related genes during etiolation and upon illumination in pea and oat. Relative transcript levels of *psaA*, *psaB*, *atpβ*, and *Lhca2* were determined by qRT-PCR in pea and oat at successive stages of 10-day-long etiolation (−10d to 0d) and for 20-day-old mature plants growing in day/night conditions. Expression levels were normalised to the geometric mean of species-specific reference genes (PP2A and TUBB for pea; Arf1 and HNR for oat). Values represent means ± SD of 3 independent biological replicates. d, with “-” days of etiolation prior to illumination or without indicating days of day/night growth; 0d, 10-day-old etiolated seedlings. Asterisks indicate statistically significant differences compared with the −10 days before light exposure (p < 0.05).

Immunoblot analysis showed PsaA at all examined stages of etiolation in pea, with a distinct band at approximately 55 kDa (Fig. 4). A faint PsaB signal was also detected at a similar molecular weight. An additional band at approximately 80 kDa was observed, the identity of which remains to be determined. PsaA abundance was substantially higher than that of PsaB throughout etiolation, with an estimated ∼2.8-fold difference when normalized to signals from mature chloroplasts; however, given that PsaB signals were close to the detection limit, this ratio should be interpreted with caution.

**Fig. 4.**
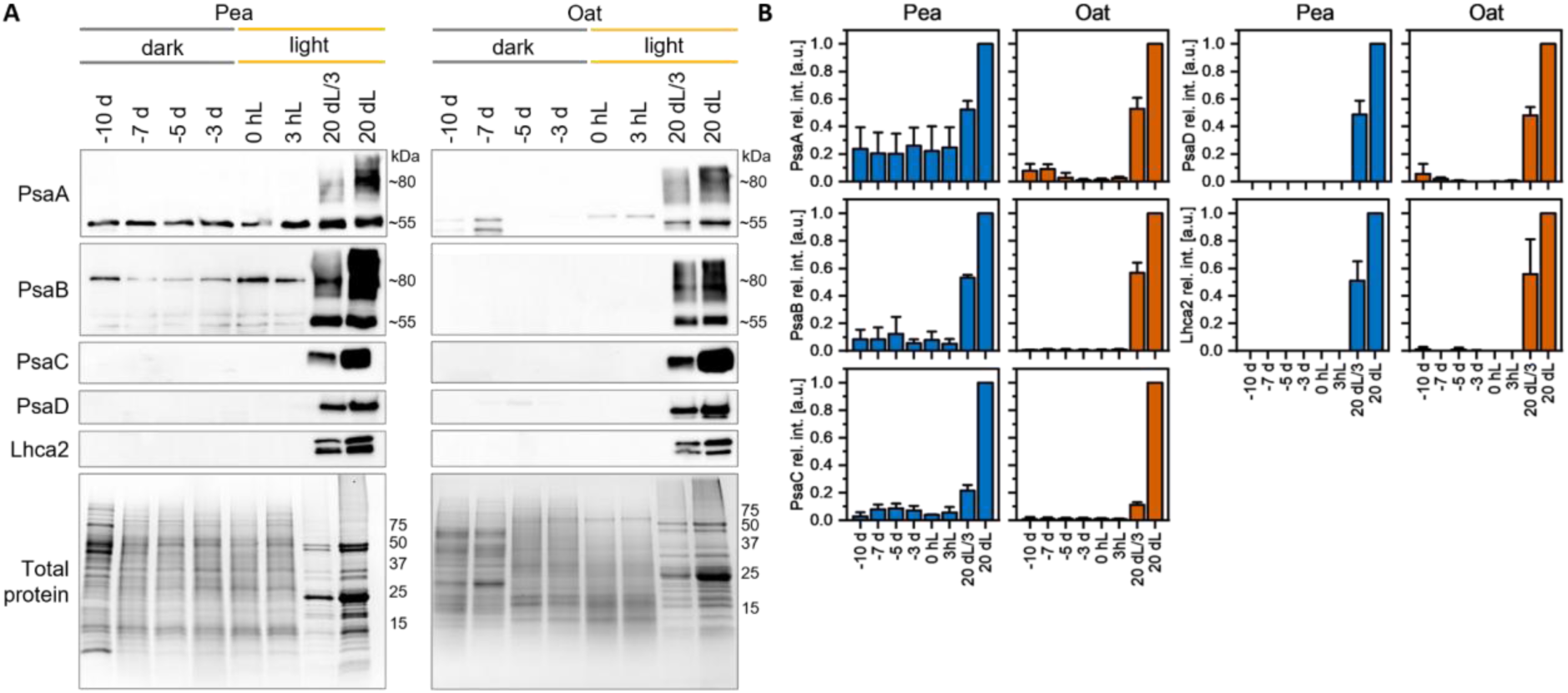
Time-course immunoblot analysis of PSI subunit accumulation during etiolation in pea and oat. (A) Total protein extracts from pea and oat seedlings harvested at successive stages of etiolation (−10d to 0d) and from mature 20-day-old day/night-grown plants were separated by SDS–PAGE and analysed by immunoblotting with antibodies against PsaA, PsaB, PsaC, PsaD, and Lhca2; representative blots from 3 independent biological replicates are shown. Equal loading was confirmed by Coomassie staining. (B) Signal intensities were quantified using ImageLab software version 6.0.1 (Bio-Rad) (Taylor *et al*., 2013) and normalized to the corresponding signal in mature samples of respective species. d, with “-” days of etiolation prior to illumination or without indicating days of day/night growth; 0d, 10-day-old etiolated seedlings.

In oat, PsaA was detected only after prolonged etiolation (10 days; 0d) as a faint band migrating at an anomalously high apparent molecular weight relative to the mature protein (Fig. 4), with a signal visible at day 3 (-7d). The intensity of the PsaA signal in oat after 10 days of etiolation was approximately 80-fold lower than in mature chloroplasts. PsaB remained below the detection limit in oat, and PsaC and PsaD were not detected at any etiolation stage in either species (Fig. 4). Lhca2, used as a light-dependent control, was not detected in either species during etiolation, consistent with its known requirement for light-driven synthesis.

These data show that PsaA accumulation in the dark is decoupled from transcript levels and occurs to a greater extent in the dicot pea than in the monocot oat. However, in neither species is it accompanied by the other PSI core or light-harvesting proteins tested.

### Blue-native PAGE separation of membrane proteins in pea etioplasts

To investigate the potential native complex associations of PsaA in etioplasts, blue-native PAGE followed by immunoblotting was performed on pea samples after 10 days of etiolation (Fig. 5). The strongest PsaA signal was detected in the low-molecular-mass region. PsaA was also detected in higher-molecular-mass regions, where it co-migrated with chlorophyll synthase, ferredoxin–NADP⁺ reductase (FNR), PsaB, and LPOR, suggesting co-localization within a shared membrane microdomain.

**Fig. 5.**
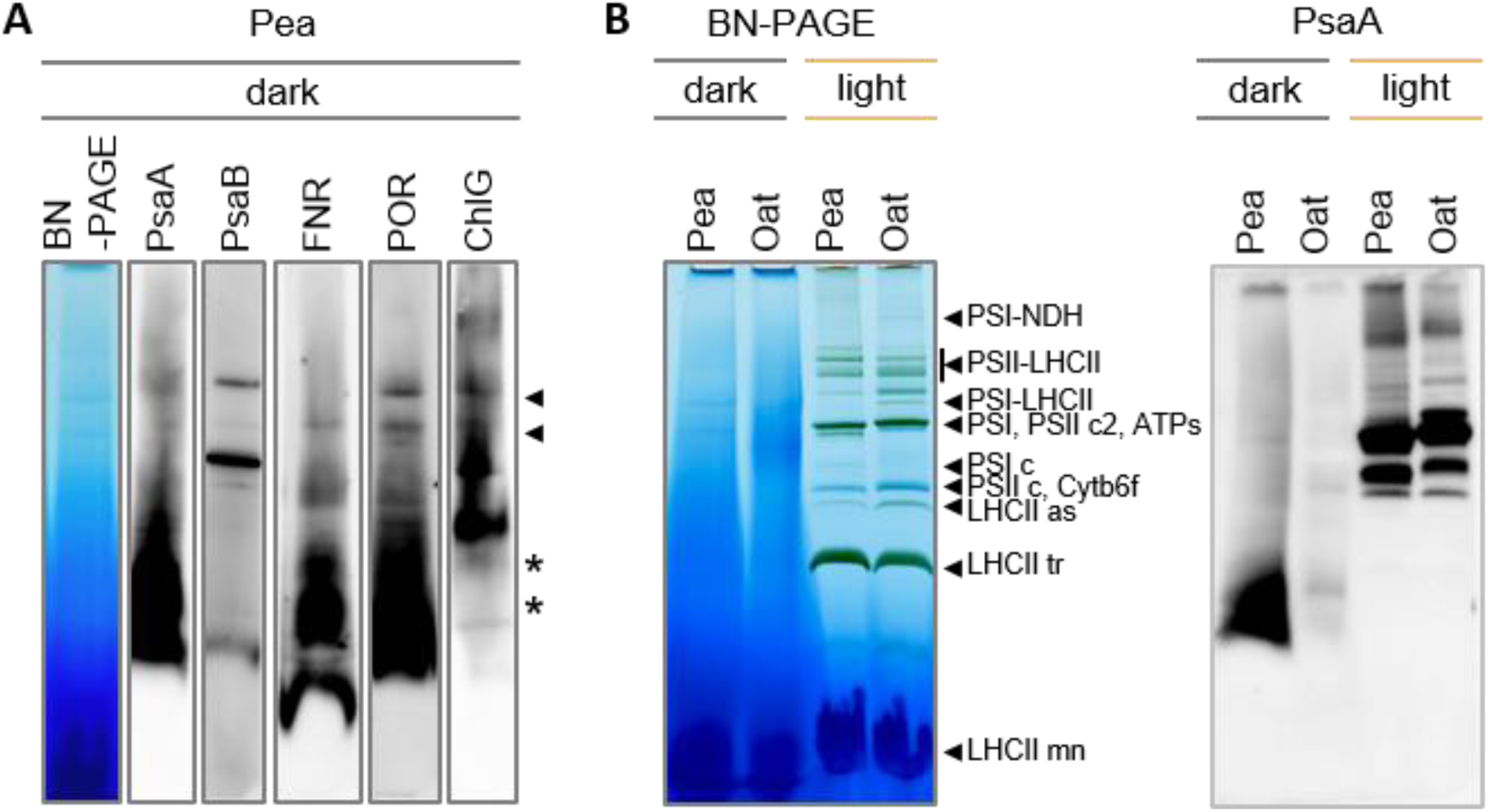
Blue-native PAGE analysis of PsaA-containing complexes in pea etioplasts. (A) Blue-native PAGE electrophoresis of pea etioplast protein complexes and corresponding immunoblotting with antibodies against PsaA, PsaB, ferredoxin–NADP⁺ reductase (FNR), POR, and chlorophyll synthase (ChlG), indicating colocalization of PsaA with these proteins in high molecular weight bands (marked with arrowhead). The low-molecular-mass region containing the predominant PsaA signal is indicated by an asterisk. (B) Etioplast membrane protein complexes of pea and oat after 10 days of etiolation and from mature 20-day-old chloroplasts were separated by blue-native PAGE and analysed by immunoblotting with an antibody against PsaA. Representative blots from 3 independent biological replicates are shown.

### Seedling de-etiolation dynamics

Finally, we set out to test whether the presence of PsaA in etiolated pea, and its co-migration with proteins involved in chlorophyll synthesis on Blue Native PAGE, could result in an accelerated onset of PSI accumulation and functionality in pea compared with oat. We used two complementary methods across the first 3 h of illumination of etiolated seedlings: low-temperature chlorophyll fluorescence and P700 signal registration (Fig. 6). The former reports on PSI presence, while the latter assesses its functionality. Chlorophyll emission spectra of illuminated seedlings, recorded at 0.5 h intervals, showed one or two dominant peaks depending on the duration of light exposure (Fig. 6A). The peak with a maximum at ∼683 nm corresponds to chlorophyll bound to PSII proteins, while the peak with a maximum above 700 nm is associated with PSI-bound chlorophyll (Banaś *et al*., 2024). The shift of these peaks at successive time points indicated complex formation and maturation, including core complex biogenesis and binding of the LHCII and LHCI antennae to PSII and PSI, respectively. The PSI-related signal was clearly visible in pea already after 1.5 h of illumination, whereas in oat, only a weak signal in this region was observed at this time point. In the following hours, however, the PSI-associated signal in pea increased only slightly, while in oat the increase was more pronounced. The low-temperature fluorescence results agree with the P700 measurements reflecting PSI functionality: the first signs of P700 oxidation were registered in pea already after 1.5 h of illumination, whereas in oat the onset of the P700 signal was delayed by 30 min (Fig. 6B). Interestingly, at later stages of illumination, we observed no differences between the two species in the course of the recorded traces. Together, the two methods reveal earlier emergence of PSI-related signals and P700 activity in pea than in oat during the first hours of illumination, with the two species converging at later stages.

**Fig. 6.**
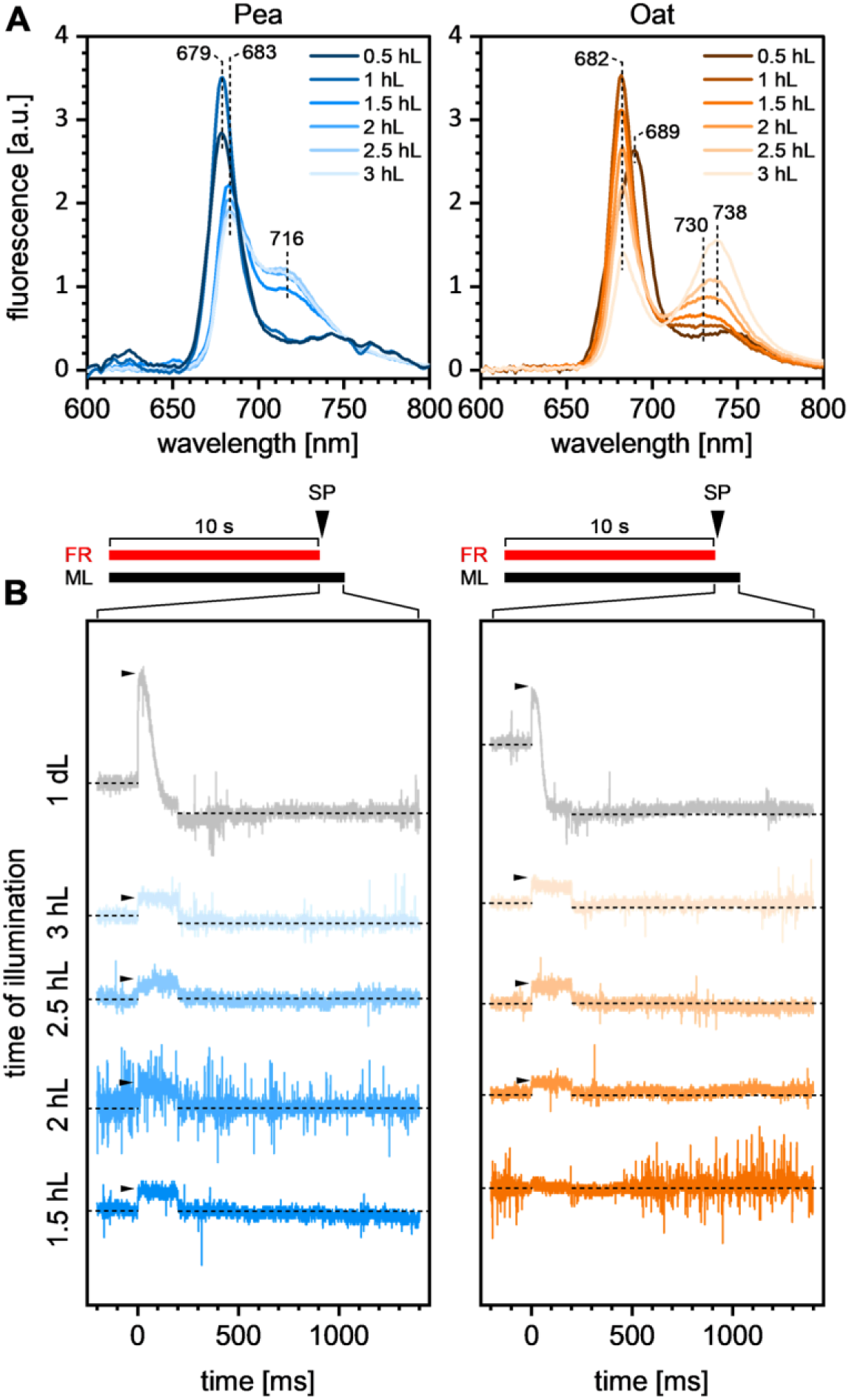
Low-temperature fluorescence spectroscopy and P700 measurements during the de-etiolation process in pea and oat. (A) Representative 77 K fluorescence emission spectra recorded from pea and oat seedlings at successive stages of light illumination (0.5 h to 3 h). Excitation wavelength: 440 nm. Emission bands at ∼680 nm correspond to chlorophyll bound with PSII, and bands with the maxima above 700 nm to chlorophyll bound with PSI. Spectra were normalized to the equal area under the curve. (B) Representative traces for the P700 signal. The increase in the signal intensity depicts P700 oxidation. FR - Far-red light; SP - Saturation Pulse; ML - Measuring Light. Results representative of 3 independent biological replicates.

## Discussion

### PsaA accumulates in etioplasts prior to illumination in a species-dependent manner

The biogenesis of PSI in higher plants is a highly coordinated process requiring the integration of chloroplast- and nuclear-encoded components (Schöttler *et al*., 2011). While the assembly and regulation of PSI have been extensively studied in mature chloroplasts (Rolo *et al*., 2024b), the possible early stages of PSI biogenesis in dark-grown seedlings remain poorly understood. In this study, we provide evidence that components of the PSI core, particularly PsaA, accumulate during etiolation in a species-dependent manner, suggesting that elements of PSI assembly may be initiated prior to light exposure. The identity of PsaA in etiolated pea was independently confirmed by MS/MS analysis, ruling out non-specific antibody cross-reactivity as an explanation for the immunoblot signal.

Our data demonstrate that PsaA is detectable in etiolated seedlings of pea from the earliest stages of development, whereas its accumulation in oat is delayed and occurs at substantially lower levels. The anomalously high apparent molecular weight of PsaA in oat etioplasts, reproducible across independent replicates and absent in mature oat chloroplasts, requires explanation. The shift could reflect SDS-resistant aggregation or stable association with another protein not fully dissociated under the denaturing conditions used. Since the synthesis of PsaA and PsaB in dark-grown seedlings has been proposed to be limited at the stage of polypeptide elongation (Klein *et al*., 1988), the higher-molecular-weight form observed in oat may additionally represent PSI intermediates that accumulate. We cannot also entirely exclude antibody cross-reactivity with a distinct protein of similar epitope, though the absence of this signal in mature oat chloroplasts argues against this interpretation.

The variability in the accumulation of PsaA is consistent with previous reports describing inconsistent detection of PSI core subunits in etioplasts across species and experimental systems (Eichacker *et al*., 1990; Kanervo *et al*., 2008; Klein *et al*., 1988; Laing *et al*., 1988; Li *et al*., 2020; Machold and Høyer-Hansen, 1976; Nechushtai and Nelson, 1985; Rudowska *et al*., 2012; Takabe *et al*., 1986; Vierling and Alberte, 1983; Yang *et al*., 2016). In vitro studies on barley etioplasts showed that *psaA/psaB* mRNAs are associated with membrane-bound polysomes in etioplasts and that translational elongation is constrained by membrane-associated factors, but can be modulated by changes in membrane integrity (Klein *et al*., 1988). The observed variability in PsaA accumulation may also be consistent with recent studies demonstrating that PSI biogenesis and PsaA/PsaB translation are sensitive to the stromal redox state and NADP(H)-dependent regulatory pathways (Bhattacharjee, 2013; Ji *et al*., 2022).

### An imbalance in PsaA and PsaB accumulation suggests that the autoregulatory model may not fully apply during early angiosperm seedling development

PsaB protein remained at low or undetectable levels in both species, indicating an imbalance in the accumulation of the two reaction center subunits during etiolation. This finding is notable in the context of proposed autoregulatory mechanisms, in which the synthesis of PsaA is thought to depend on the prior presence of PsaB. This relationship was established primarily in Chlamydomonas and not yet fully tested during angiosperm ontogenesis (Stampacchia *et al*., 1997; Wostrikoff *et al*., 2004). While PsaB was detected at low levels near the detection limit in pea etioplasts, its relative underrepresentation compared to PsaA raises the possibility that this regulatory coupling may operate differently (or with different kinetics) during early seedling development in angiosperms. Importantly, recent evidence indicates that this hierarchical, Control by Epistasy of Synthesis-like translational feedback is not strictly conserved in land plants, where the synthesis of PSI core subunits appears to be less tightly coupled to PSI assembly than in Chlamydomonas (Ghandour *et al*., 2025), supporting the idea that PsaA and PsaB accumulation in higher plants may be differentially regulated during ontogenesis. We cannot exclude, however, that PsaB is present at levels below reliable detection, and a more definitive assessment would require more sensitive quantitative approaches.

The presence of PsaA at early stages of etiolation may reflect either *de novo* synthesis or the persistence of plastid-derived proteins from embryonic development. While our data do not distinguish between these possibilities, the consistent detection of PsaA across multiple time points argues for active maintenance or synthesis rather than transient carryover alone. Despite the presence of PsaA, other PSI core subunits, including PsaC and PsaD, were not detected under etiolation conditions. Together with the low abundance of PsaB, this suggests that PSI assembly does not progress beyond early stages in darkness. Instead, PsaA appears to accumulate in isolation or in association with non-PSI protein complexes.

### PsaA co-localizes with chlorophyll biosynthesis machinery in membrane microdomains

Blue-native PAGE co-migration of PsaA with LPOR, chlorophyll synthase, and FNR in higher-molecular-mass fractions is notable because BN-PAGE employs mild detergent conditions that preserve native membrane complexes and microdomain organization. Co-migration under these conditions therefore suggests these proteins may localize in a shared membrane-associated environment of membrane regions within the prothylakoid or PLB system of etioplasts, consistent with the organized structural and functional differentiation of developing plastid membranes (Floris and Kühlbrandt, 2021). In higher plants, thylakoid membranes are known to exhibit pronounced lateral heterogeneity, giving rise to spatially segregated protein complexes, which support photosynthetic function and the spatial organization of photosynthetic complex assembly (Trotta *et al*., 2025). Additionally, microdomain organization has been described in thylakoids and implicated in the spatial coordination of photosynthetic protein biogenesis in Chlamydomonas, where distinct membrane domains facilitate the stepwise assembly of membrane complexes (Schottkowski *et al*., 2012; Sun *et al*., 2025). The possible co-localization of PsaA with LPOR, FNR and chlorophyll synthase within the same membrane proximity may therefore reflect a spatially organized biosynthetic environment in which chlorophyll precursors and nascent PSI apoproteins are in close proximity, potentially facilitating efficient pigment loading upon illumination.

In this context, the organization of LPOR within the PLB has been proposed to reflect a pre-organized etioplast architecture that enables efficient light-triggered conversion of Pchlide to chlorophyllide, followed by diffusion and rapid utilization of chlorophyll intermediates within the membrane environment (Floris and Kühlbrandt, 2021), which may provide a general contextual framework for considering potential spatial relationships between chlorophyll biosynthesis and PSI apoprotein assembly during early chloroplast development.

### Etioplasts may provide a preparatory environment for PSI assembly upon illumination

The elevated levels of Alb3 in etioplasts, particularly in dicot species, together with the pre-illumination accumulation of PsaA, suggest that at least some molecular prerequisites for PSI biogenesis – including the factors required for membrane protein insertion (Pasch *et al*., 2005) – are established prior to light exposure.

However, it should be noted that Alb3 is a protein involved in the insertion of multiple thylakoid membrane complex subunits, as it has been shown to interact not only with PsaA but also with components of e.g., PSII and the ATP synthase complex (Pasch *et al*., 2005; Woolhead *et al*., 2001). Importantly, the higher Alb3 abundance in dicot etioplasts compared with oat may directly contribute to the greater accumulation of PsaA observed in dicots. If Alb3-mediated membrane insertion is a rate-limiting step in early PsaA integration, then its elevated levels in dicot etioplasts could facilitate co-translational or post-translational insertion of PsaA into the etioplast membrane even in the absence of light. The comparatively lower Alb3 levels in oat etioplasts would, on this view, represent one component of a broader attenuation of PSI biogenesis in monocots prior to illumination, however, poor reactivity of Alb3 antibody in oat cannot be excluded.

Although testing the rapid, light-driven process of PSI assembly is a complex task beyond the scope of this work, we showed that the presence of PsaA in pea in darkness corresponds with a more rapid appearance of signals reflecting PSI accumulation and functionality in this species compared with oat, in which the process is delayed by 30 min. These results suggest that PsaA accumulated in darkness, in close membrane proximity to the proteins responsible for light-dependent chlorophyll synthesis, can be utilized during the initial onset of photosynthesis. Interestingly, after further hours of illumination, the oat seedlings rapidly catch up in PSI accumulation, and the increase in the PSI-related signal in the low-temperature spectra is more pronounced than in pea, reflecting the higher level of *psaA* transcript in oat at the end of the night period compared with the beginning of etiolation. In light of these results, the dark accumulation of PsaA appears to be an important step toward establishing autotrophy in pea, giving the seedling a pool of this protein ready for assembly at the very onset of illumination. Without this nighttime build-up, PSI formation would depend on synthesizing PsaA *de novo* from a low transcript pool, which could substantially retard the process in pea.

### Monocots and dicots differ in the balance between pigment priming and PSI apoprotein accumulation in darkness

A notable finding of this study is the pronounced difference between pea and oat in both PsaA accumulation and pigment dynamics. While pea maintains relatively stable levels of Pchlide complexes and exhibits early PsaA accumulation, oat shows a strong increase in the Pchlide:LPOR:NADPH ratio over time but only minimal accumulation of PsaA. In oat, efficient accumulation of the Pchlide:LPOR:NADPH complex may prime the system for rapid chlorophyll production upon illumination, whereas PSI core protein synthesis remains tightly restricted. In contrast, pea appears to permit partial accumulation of PSI apoproteins in advance of pigment availability. Such variation may reflect differences in developmental programs between monocots and dicots, or species-specific regulatory mechanisms operating at multiple levels of chloroplast biogenesis in darkness. These findings may help explain previously reported inconsistencies in the detection of PSI components in etioplast-related proteomic analyses – particularly the reported absence (Klein *et al*., 1988; Laing *et al*., 1988; Machold and Høyer-Hansen, 1976; Takabe *et al*., 1986; Vierling and Alberte, 1983), or low abundance of PsaA in monocot plants (Nechushtai and Nelson, 1985) and underscore the importance of considering species context in studies of plastid development.

### Dark accumulation of PsaA may represent an early light-independent step in PSI biogenesis

Current models of PSI biogenesis emphasize the requirement for light, chlorophyll availability, and coordinated synthesis of PsaA and PsaB (Komenda and Sobotka, 2019; Yang *et al*., 2015), although recent findings suggest that the hierarchical coordination of PSI core subunit synthesis may be less stringent in higher plants than previously assumed (Ghandour *et al*., 2025). We demonstrated that at least one core subunit, PsaA, can accumulate in darkness in certain species, suggesting that early steps of PSI assembly, including membrane insertion of PsaA, may occur independently of light-driven processes. However, the absence of downstream subunits and functional complexes indicates that full PSI assembly remains tightly regulated and likely requires additional light-dependent signals, such as chlorophyll availability or redox cues. The accumulation of PsaA in darkness may therefore represent an early light-independent step in PSI biogenesis, suggesting that the boundary between skotomorphogenic and photomorphogenic development is less discrete than current models imply.

## Supplementary data

Supplementary materials include Tab. 1 with primers used in this study and Tab. 2 with a list of proteins identified in the etioplasts fraction isolated from 10-day etiolated (0hL) pea seedlings.

## Author contributions

AW and ŁK conceived the study; AW, ŁK, RM, KW, and PG planned and designed the research; AW, KW, RM, KG performed experiments; AW, RM, KG, PG, ŁK analyzed data; AW wrote the original draft; all authors reviewed the draft and approved the final version.

## Conflict of interest

The authors declare no conflict of interest.

## Funding

This research was funded by the National Science Centre, Poland grant PRELUDIUM 2023/49/N/NZ3/01212.

## Data availability

The data that support the findings of this study are openly available in Dane Badawcze UW repository at doi:10.58132/P6WDSS reference number

## Notes

### Competing Interest Statement

The authors have declared no competing interest.

